# *P. aeruginosa* tRNA-fMet halves secreted in outer membrane vesicles suppress lung inflammation in Cystic Fibrosis

**DOI:** 10.1101/2024.02.03.578737

**Authors:** Zhongyou Li, Roxanna Barnaby, Amanda Nymon, Carolyn Roche, Katja Koeppen, Alix Ashare, Deborah A. Hogan, Scott A. Gerber, Douglas J. Taatjes, Thomas H. Hampton, Bruce A. Stanton

## Abstract

Although tobramycin increases lung function in people with cystic fibrosis (pwCF), the density of *Pseudomonas aeruginosa (P. aeruginosa)* in the lungs is only modestly reduced by tobramycin; hence, the mechanism whereby tobramycin improves lung function is not completely understood. Here, we demonstrate that tobramycin increases 5′ tRNA-fMet halves in outer membrane vesicles (OMVs) secreted by laboratory and CF clinical isolates of *P. aeruginosa*. The 5′ tRNA-fMet halves are transferred from OMVs into primary CF human bronchial epithelial cells (CF-HBEC), decreasing OMV-induced IL-8 and IP-10 secretion. In mouse lung, increased expression of the 5′ tRNA-fMet halves in OMVs attenuated KC secretion and neutrophil recruitment. Furthermore, there was less IL-8 and neutrophils in bronchoalveolar lavage fluid isolated from pwCF during the period of exposure to tobramycin versus the period off tobramycin. In conclusion, we have shown in mice and *in vitro* studies on CF-HBEC that tobramycin reduces inflammation by increasing 5′ tRNA-fMet halves in OMVs that are delivered to CF-HBEC and reduce IL-8 and neutrophilic airway inflammation. This effect is predicted to improve lung function in pwCF receiving tobramycin for *P. aeruginosa* infection.

**New and noteworthy:** The experiments in this report identify a novel mechanim whereby tobramycin reduces inflammation in two models of CF. Tobramycin increased the secretion of tRNA-fMet haves in OMVs secreted by *P. aeruginiosa*, which reduced the OMV-LPS induced inflammatory response in primary cultures of CF-HBEC and in mouse lung, an effect predicted to reduce lung damage in pwCF.

**Graphical abstract:** The anti-inflammatory effect of tobramycin mediated by 5′ tRNA-fMet halves secreted in *P. aeruginosa* OMVs. **(A)** *P. aeruginosa* colonizes the CF lungs and secrets OMVs. OMVs diffuse through the mucus layer overlying bronchial epithelial cells and induce IL-8 secretion, which recruits neutrophils that causes lung damage. (**B**) Tobramycin increases 5′ tRNA-fMet halves in OMVs secreted by *P. aeruginosa*. 5′ tRNA-fMet halves are delivered into host cells after OMVs fuse with lipid rafts in CF-HBEC and down-regulate protein expression of MAPK10, IKBKG, and EP300, which suppresses IL-8 secretion and neutrophils in the lungs. A reduction in neutrophils in CF BALF is predicted to improve lung function and decrease lung damage.

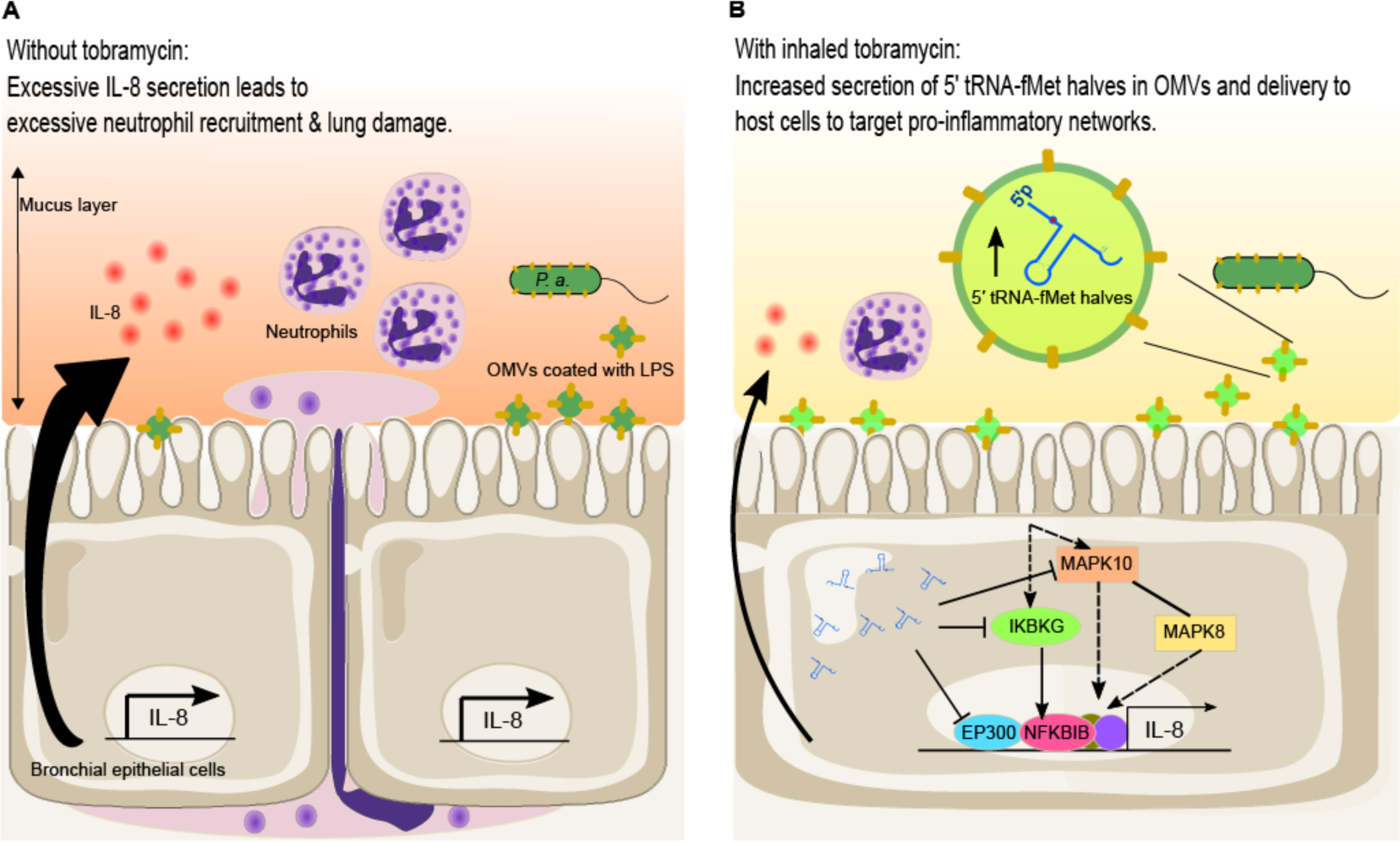

## Introduction

CF is a genetic disease caused by absent or aberrant function of the cystic fibrosis transmembrane conductance regulator (CFTR), which leads to airway periciliary dehydration, increased mucus viscosity, and decreased mucociliary clearance (1, 2). Insufficient mucociliary clearance facilitates persistent bacterial infection, non-resolving lung inflammation, and excessive neutrophil recruitment (3, 4). Chronic neutrophilic airway inflammation damages the lungs by continuous secretion of reactive oxygen species (ROS) and proteases, contributing to bronchiectasis and progressive lung function loss in people with CF (pwCF) (5, 6). Although highly effective modulator therapy (HEMT) improves lung function, and decreases hospitalization in pwCF, HEMT does not eliminate *P. aeruginosa* lung infections or the hyperinflammatory response to chronic infections (7–9).

*Pseudomonas aeruginosa* (*P. aeruginosa*) is an opportunistic pathogen that infects the lungs of immunocompromised individuals, including those with chronic obstructive pulmonary disease and cystic fibrosis (CF), and is an important cause of acute and ventilator-associated pneumonia (10–14). *P. aeruginosa* chronically colonizes the lungs of ∼50% of adults with CF, and its presence is strongly associated with reduced forced expiratory volume (FEV_1_) and a progressive loss of lung function (1, 7, 15). Inhaled tobramycin is the most commonly used antibiotic to suppress *P. aeruginosa* infections in pwCF and to ameliorate lung function loss once chronic pulmonary colonization is established (7, 16). The long-term use of inhaled tobramycin significantly improves lung function and reduces mortality in pwCF (17, 18). Inhaled tobramycin is administered in intermittent repeated cycles of 28 days on the drug and 28 days off. In a double-blind, placebo-controlled study, lung function improved significantly after the first two weeks of treatment and correlated with a decrease of *P. aeruginosa* colony-forming units (CFUs) in sputum by more than 158-fold (19). Intriguingly, the magnitude of the reduction in bacterial CFUs was less than 10-fold in the third cycle of therapy, although lung function improvement was maintained at a comparable level (19). Furthermore, an open-label, follow-on trial with adolescent patients and 12 treatment cycles revealed that the reduction of *P. aeruginosa* CFUs in sputum only explained 11.7% of CF lung function improvement (20). Moreover, a more recent analysis of sputum in pwCF revealed that one cycle of tobramycin reduced *P. aeruginosa* abundance by only ∼1 log after one week (21). Together, these data suggest that tobramycin improves CF lung function by an unknown mechanism in addition to its bactericidal activity. Accordingly, the goal of this study was to identify non-bactericidal effects of tobramycin.

In the CF lungs, *P. aeruginosa* resides primarily in the mucus overlying lung epithelial cells and secretes outer membrane vesicles (OMVs), 50-300 nm lipopolysaccharide (LPS)-decorated vesicles secreted by all Gram-negative bacteria (22–24), that fuse with and deliver their content to lung epithelial cells and stimulate the immune response by human bronchial epithelial cells (HBEC) (22, 25–27). We and others have shown that secreted OMVs stimulate the inflammatory response in the lungs and also fuse with lipid rafts in host cells and deliver virulence factors, DNA, small RNAs (sRNAs), and transfer RNA (tRNA) fragments into HBEC, which mitigate the host immune response to OMVs (22, 25–29). For example, *P. aeruginosa* secretes a 24-nucleotide (nt) small RNA (sRNA) in OMVs, which diffuse through the airway mucus layer, fuse with lipid rafts in HBEC and transfer the sRNA into HBEC, which inhibits IL-8 secretion (30, 31). The sRNA also mitigates the OMV-induced secretion of IL-8 (KC in mice), a potent neutrophil attractant, leading to attenuated recruitment of neutrophils into mouse lungs (30).

This study aimed to test the hypothesis that tobramycin prevents the decline in lung function of pwCF, at least in part, by increasing the level of anti-inflammatory sRNAs in OMVs secreted by *P. aeruginosa.* Here, we demonstrate that tobramycin increases the abundance of two 5′ formyl-methionine tRNA halves (tRNA-fMet halves) in OMVs, and that the tRNA-fMet halves are delivered into primary CF-HBECs by OMVs. The tRNA-fMet halves suppress the OMV-stimulated increase in IL-8 and IP-10 secretion by CF-HBECs. Moreover, OMVs secreted by tobramycin treated *P. aeruginosa* reduced KC (a murine homolog of IL-8) levels and neutrophils in mouse lungs compared to OMVs secreted by vehicle exposed *P. aeruginosa*. We also report that in pwCF the IL-8 concentration and neutrophil content in bronchoalveolar lavage fluid (BALF) is significantly reduced during the month on tobramycin compared to the month off tobramycin. Taken together, these data suggest that the clinical benefit of tobramycin is due in part to an increase in the secretion of tRNA-fMet halves in OMVs, leading to attenuation of *P. aeruginosa* IL-8 and neutrophil-mediated CF lung damage.

## Methods

### P. aeruginosa cultures

*P. aeruginosa* (strain PA14) and all CF clinical isolates were grown in luria broth (LB, ThermoFisher Scientific, Waltham, MA) liquid cultures at 37°C with shaking at 225 rpm for 16 hours. Tobramycin (1 μg/mL), a concentration that reduces *P. aeruginosa* by an amount similar to that observed clinically (19–21), or vehicle was added to the cultures. The CF clinical isolates of *P. aeruginosa*, two mucoid and two non-mucoid strains, have been characterized previously (32, 33). In some experiments, the 5′ tRNA-fMet1 half (5′-CGCGGGGTGGAGCAGTCTGGTAGCTCGTCGGGCTC-3′) was cloned into the arabinose-inducible expression vector pMQ70 (34) by cutting EcoRI and SmaI restriction sites. GenScript (GenScript USA Inc., Piscataway, NJ, USA) performed the cloning procedure. PA14 was transformed with the 5′ tRNA-fMet1 half expression vector or empty vector via electroporation. *P. aeruginosa* strains with the arabinose-inducible vector and its derivatives were grown in LB with 133 mM L-arabinose (2% w/v) and 300 μg/ml carbenicillin (both from Sigma-Aldrich).

### Growth kinetics of *P. aeruginosa*

*P. aeruginosa* overnight cultures in LB were centrifuged, washed, and resuspended in fresh LB before measuring the optical density at 600 nm (OD600) to determine cell number. Bacteria were seeded at 1×10^5^ cells per 100 µl LB with or without tobramycin (1 μg/mL) in a transparent, flat bottom, 96-well plate covered with a lid. The plate was cultured in a plate reader at 37°C for 24 h. The reader was programmed to measure the OD600 every 15 minutes after shaking the plate for 5 seconds.

### LIVE/DEAD BacLight Bacterial Viability Assay

The LIVE/DEAD viability assay was performed on *P. aeruginosa* exposed to vehicle or tobramycin (1 μg/ml) according to the manufactures protocol (ThermoFisher Scientific).

### Outer membrane vesicle preparation and quantification

OMVs were isolated and characterized as described by us previously (35, 36). Briefly, *P. aeruginosa* overnight cultures were centrifuged for 1 h at 2800 g and 4°C to pellet the bacteria. The supernatant was filtered twice through 0.45 μm PVDF membrane filters (Millipore, Billerica, MA, USA) to remove bacteria (confirmed by colony counts) and concentrated with 30K Amicon filters (Millipore, Billerica, MA, USA) at 2800 g and 4°C to obtain ∼200 μL concentrate. The concentrate was resuspended in OMV buffer (20 mM HEPES, 500 mM NaCl, pH 7.4) and subjected to ultracentrifugation for 2 h at 39,000 g (21,000 RPM) and 4°C to pellet OMVs. OMV pellets were re-suspended in 60% OptiPrep Density Gradient Medium (Sigma-Aldrich, Cat. # D1556) and layered with 40%, 35%, 30% and 20% OptiPrep diluted in OMV buffer. OMVs in OptiPrep layers were centrifuged for 16 h at 100,000 g (31,00 RPM) and 4°C. 500 μl fractions were taken from the top of the gradient, with OMVs residing in fractions 2-3, corresponding to 25% OptiPrep. The purified OMVs were quantified by nanoparticle tracking analysis (NTA, NanoSight NS300, Malvern Panalytical Ltd, Malvern, UK) and negative staining electron microscopy before exposure of CF-HBECs or mice to OMVs.

### CF-HBEC culture

De-identified primary human bronchial epithelial cells from six CF donors (CF-HBECs, Phe508del homozygous) were obtained from Dr. Scott Randell (University of North Carolina, Chapel Hill, NC, USA) and cultured as described previously (36, 37). Since the cells were acquired from discarded tissue, and comprised no patient identifiers, their use in this study was not considered as human subject research by the Dartmouth Committee for the Protection of Human Subjects. Briefly, cells were grown in BronchiaLife basal medium (Lifeline Cell Technology, Frederick, MD, USA) supplemented with the BronchiaLife B/T LifeFactors Kit (Lifeline) as well as 10,000 U/ml Penicillin and 10,000 μg/ml Streptomycin. The cultures were determined to be sterile and free of mycoplasma contamination.

To polarize cells, CF-HBECs were seeded on polyester transwell permeable filters (#3412 for 24-mm Transwell Corning, Corning, NY) coated with 50 μg/ml Collagen type IV (Sigma-Aldrich, St. Louis, MO). Air Liquid Interface (ALI) medium was added to both apical and basolateral sides for plating. 24-48 hours after plating the apical media was removed and cells were cultured at ALI and fed basolaterally every other day with ALI media for 3-4 weeks before cells were fully polarized for treatment (38).The effect of OMVs on CF-HBEC viability was assessed using the CytotTox96 Non-Radio. Cytotoxicity Assay™(Promega Catalog number G1781) according to the manufactures recommendations. There was no effect of OMVs on cytotoxicity (n=4 donors).

### Exposure of CF-HBEC to OMVs

Polarized CF-HBEC on 24-mm Transwell filters were gently washed with PBS, and 1.5 mL of ALI medium was added to the basolateral side. 6.0 ×10^10^ OMVs or the same volume of vehicle control media run throught the OMV isolation step were applied to the apical side of cells. 8.4 ×10^10^ Tobi-OMVs (1.4X Tobi-OMVs) were also used. OMVs were added to the apical side in a total of 500ul ALI medium. Previous studies have shown that the concentration of OMVs in biological fluids including BALF are ∼10^10^ /ml (reviewed in (28, 29)). After a six-hour exposure, the basolateral media was collected for cytokine measurements.

### Cytokine measurements

Cytokine secretion in CF BALF was measured with the Human IL-8/CXCL8 DuoSet ELISA (#DY208, R&D Systems, Minneapolis, MN). KC in mouse lungs was analyzed with the Mouse CXCL1/KC DuoSet ELISA (#DY453, R&D Systems, Minneapolis, MN). Cytokines secreted by primary CF-HBECs were analyzed by the MILLIPLEX MAP Human Cytokine/Chemokine 48-Plex cytokine assay (Millipore). Cytokines were measure in the basolateral solution only since CF-HBEC were grown in air-liquid interface culture and there was insufficient apical fluid to measure cytokines.

### RNA isolation and small RNA-seq analysis

PA14 was grown in T-broth lacking yeast (10 g tryptone and 5 g NaCl in 1 L H_2_O) with or without tobramycin (1 μg/mL) to reduce small RNA reads from yeast present in LB medium. The culture supernatants were processed as mentioned above to obtain OMV pellets. The pellets were resuspended with OMV buffer and re-pelleted again by centrifugation at 100,000 g (31,000 RPM) for 2 h at 4°C and lysed with Qiazol followed by RNA isolation with the miRNeasy kit (Qiagen, Germantown, MD) to obtain total RNA including the small RNA fraction. DNase-treated total RNA was used to prepare cDNA libraries with the SMARTer smRNA-Seq Kit (Takara Bio, Mountain View, CA). Libraries were sequenced as 50 bp single-end reads on an Illumina HiSeq sequencer. The first three nucleotides of all reads and the adapter sequences were trimmed using cutadapt (39) before sequence alignment.

To verify the overexpression of 5′ tRNA-fMet1 half, PA14 clones with the 5′ tRNA-fMet1 half expression plasmid or the empty pMQ70 vector were grown in LB (with L-arabinose and carbenicillin) for isolation of EV-OMVs and tRNA1-OMVs. The OMV pellets were collected and processed as described above to isolate RNA. DNase-treated total RNA was further treated with RNA 5’ Pyrophosphohydrolase (New England Biolabs) to remove possible pyrophosphate from the 5’ end. The QIAseq miRNA Library Kit (Qiagen) was used to prepare cDNA libraries, and 50 bp single-end sequencing was performed on an Illumina MiniSeq system.

Reads were aligned to the PA14 reference genome using CLC Genomics Workbench (CLC-Bio/Qiagen) with the following modifications from the standard parameters: a) the maximum number of mismatches = zero to eliminate unspecific alignment and b) the maximum number of hits for a read = 30 to capture all sRNAs aligned to the PA14 genome. Pileups of mapped reads and frequency tables for each unique sequence were exported for normalization and further analysis with the software package edgeR in the R environment (40, 41).

### Detection of 5′ tRNA-fMet halves by RT-PCR

The induction of 5′ tRNA-fMet halves by tobramycin in OMVs secreted by PA14 and four *P. aeruginosa* clinical strains was detected by custom Taqman Small RNA Assay (#4398987, ThermoFisher Scientific), according to the manufacturer′ s instructions. cDNA was synthesized with the TaqMan MicroRNA Reverse Transcription Kit (#4366596, ThermoFisher Scientific). PCR amplification and detection of 5′ tRNA-fMet halves were performed using the TaqMan Universal PCR Master Mix (#4304437, ThermoFisher Scientific) as well as custom primers and probe design to target both 5′ tRNA-fMet halves specifically.

### Electron microscopy of OMV

OMVs were imaged by negative staining transmission electron microscopy. Ten microliters of each sample were incubated for 2 minutes on a freshly glow discharged 200 mesh nickel grid, followed by wicking of the solution with filter paper, and rinsing on seven sequential drops of Millipore filtered distilled water. The still moist grids were then touched to one drop of NanoW tungsten stain (Ted Pella; product #09S432, lot #14702), the excess was wicked off immediately with filter paper before repeating this step on a second drop of Nano-W, which was left for 1 minute before being wicked off and allowed to dry. The stained grids were imaged at 80kV in a JEOL 1400 transmission electron microscope (JEOL Inc., Danvers, MA), and digital images acquired in TIFF with an AMT XR11 digital camera (AMT, Woburn, MA). For measurement of OMVs by electron microscopy, images were opened in MetaMorph Offline image analysis software (version 7.8.0.0; Molecular Devices LLC, San Jose CA), and the diameters were determined by drawing a measuring line from one side of the vesicle to the other. The average diameter and standard deviation (SD) were then calculated from these measurements. Image acquisition and EV diameter measurements were performed by an investigator blinded to the sample treatment.

### 5′ tRNA-fMet1 half target prediction

The miRanda microRNA target scanning algorithm (v3.3a) was used to predict human target genes of 5′ tRNA-fMet1 half (42). The 5′ tRNA-fMet1 half sequence was scanned against human RNA sequences (annotations from GRCh38.p13 assembly) with a mimimum miRanda alignment score of 150 to generate a list of predicted target genes and the corresponding interaction minimum free energies. To account for the effect of gene expression on target prediction, for each predicted target the minimum free energy was multiplied by the gene expression level (log2CPM) in polarized HBECs identified in our previous publication (43) to obtain an energy-expression score. 1518 genes (8.4% of all human genes) with energy-expression scores small than -200 were defined as predicted targets for the Ingenuity Pathway Analysis (44).

### Proteomic analysis

Primary CF-HBECs from 4 donors were polarized on 24 mm Transwell filters and washed with DPBS (ThermoFisher Scientific) before treatment. 2 mL ALI medium was added to the basolateral side. 2.8×10^10^ purified V-OMVs or tRNA1-OMVs in 800 μL ALI medium were applied to the apical side of cells. After a six-hour exposure, the cells were washed with DPBS and detached from the Transwell filters with pre-warmed 37°C trypsin/EDTA. Cells were pelleted and flash-frozen in liquid nitrogen for proteomic analysis. A pair of samples from one donor was excluded from the final analysis because they were accidentally mislabeled during the cell pellet preparation. The cell pellets were lysed in 8M urea/50mM Tris pH 8.1/100mM NaCl+ protease inhibitors (Roche cat # 4693116001) and quantified by BCA assay (Pierce cat # 23223), followed by trypsin digestion and desalting. 40 micrograms of peptides from each pellet were labeled with unique TMT reagent isobars; the individual TMT-labeled samples were then combined and fractionated offline into 12 fractions by PFP-RP-LC (45), followed by analysis on a UPLC-Orbitrap Fusion Lumos tribrid instrument in SPS-MS3 mode (46). The resulting tandem mass spectra were data-searched using Comet; TMT reporter ion intensities were summed for each protein and normalized for total intensity across all channels. Mean fold changes comparing tRNA1-OMV-exposed cells with EV-OMVs-exposed cells were calculated for each protein detected in all samples. Proteins were ranked by paired t-test *P* value, and network analysis of the top 20% proteins was performed with Ingenuity Pathway Analysis (IPA).

### Transfection of CF-HBECs with 5′ tRNA-fMet halves inhibitor and OMV exposure

CF-HBECs were seeded on 12-well plates (Corning Inc.) at 50,000 cells per well. Two days after seeding (∼80% confluence), cells were washed and fed with the complete Lifeline medium plus antibiotics and transfected with 50 nM custom mirVana miRNA inhibitor (inhibitor sequence: 5′-GAGCCCGACGAGCUACCAGACUGCUCCA-3′, #4464086, ThermoFisher Scientific) or 50 nM mirVanna inhibitor negative control#1 (#4464077, ThermoFisher Scientific: confirmed by ThermoFisher to have no binding sites in *P. aeruginosa* or human genes) using HiPerFect transfection reagent (Qiagen). Six hours after transfection, cells were exposed to OptiPrep vehicle control, EV-OMVs (0.4×10^10^ per well), or 1.4X Tobi-OMVs (0.55×10^10^ per well) for another 6 hrs., and then supernatants were collected for cytokine measurements.

### Mouse exposure to OMVs

All animal experiments were approved by the Dartmouth College IACUC committee (Protocol No. 00002026). 8–9 weeks old male and female C57BL/6J mice (The Jackson Laboratory, Bar Harbor, ME, USA) were inoculated by oropharyngeal aspiration with OMVs (0.5×10^10^ OMVs per mouse) or vehicle following brief anesthesia with isoflurane. OMV concentrations were adjusted with PBS to obtain 50 μl inoculation volume. 5 h after exposure, mice were euthanized using isoflurane anesthesia, followed by cervical dislocation after breathing stops. Mice tracheae were surgically exposed, and a catheter tube was inserted into the trachea and stabilized with sutures (#100–5000, Henry Schein Inc., Melville, NY, USA). The catheter was prepared by fitting a 23-gauge needle (BD #305145, Becton, Dickinson and Company, Franklin Lakes, NJ, USA) into transparent plastic tubing (BD #427411). BALF was collected by pumping 1 ml of sterile PBS into the lungs and recovered with a syringe (BD #309659). This process was repeated once to collect 2 mL of BALF.

### Human subjects and bronchoscopy

All CF subjects were enrolled in a protocol approved by the Dartmouth Health IRB (Protocol No. 22781). CF subjects (all male: three donors were homozygous for the Phe508del mutation, and one was Phe508del/type IV mutation [the Dartmouth IRB required that the second mutation be reported as Type IV to ensure subject confidentially]) prescribed with an inhaled tobramycin regimen were enrolled if they had an FEV1 > 50% predicted, and were not currently having an exacerbation. Following written informed consent, local anesthesia with nebulized lidocaine was administered to the posterior pharynx. Under conscious sedation, a flexible fiberoptic bronchoscopy was performed transorally. BALF was obtained from tertiary airways. After the bronchoscopy procedure, CF subjects were monitored per institutional protocol until they were stable for discharge.

### Quantification of neutrophils in BALF

Cells in BALF samples were pelleted and resuspended in 100 μL RBC lysis buffer (Promega) for 1 min. After removing red blood cells, the total number of cells in each BALF sample was counted, and concentrations were adjusted. 2×10^5^ cells per sample were spun onto glass slides, air-dried, and stained with the Differential Quik Stain Kit (Polysciences, Warrington, PA) according to the manufacturer′ s protocol. Neutrophils were counted under 100x magnification using a microscope. The neutrophil concentration of BALF was calculated by accounting for the retrieved BALF volume and the dilution factors used to adjust the cell concentration.

### Statistics

Data were analyzed by a variety techniques, as appropriate for each experiment and as noted in each figure legend. Data were anayzed using the R software environment for statistical computing and graphics version 4.1.0 (40) and Ingenuity Pathway Analysis (44). Statistical significance was calculated using a mixed effect linear model, Wilcoxon rank-sum tests, paired and unpaired t-tests, and likelihood ratio tests on gene-wise negative binomial generalized linear models, as indicated in the figure legends. Data were visualized, and figures were created using the R package ggplot2 (47).

### Study approval

Mouse studies were approved by the Dartmouth College IACUC (Protocol No. 00002026). Collection of BALF from pwCF was approved by the Dartmouth Health IRB committee (Protocol No. 22781) and written informed consent was received prior to participation. Since CF-HBEC were acquired from discarded tissue, and comprised no patient identifiers, their use in this study was not considered as human subject research by the Dartmouth Committee for the Protection of Human Subjects.

### Data availability

Small RNA-seq data are available from the Gene Expression Omnibus database (accession number GSE183895 and GSE183897). All other data are available from the corresponding author upon request.

### Supplemental material

Figure S1 provides validation that the 5’tRNA-fMet1 half are over-expressed in OMVs. Table S1 contains the human BALF sample collection dates.

## Results

### Tobramycin increased OMV secretion by *P. aeruginosa*

To begin to test the hypothesis that tobramycin alters the immunogenecity of *P. aeruginosa* OMVs, PA14 was exposed to vehicle or tobramycin overnight. Experiments using the LIVE/DEAD BacLight Bacterial Viability kit™ were conducted to examine the effect of tobramycin (1 μg/ml) on *P. aeruginosa* viability. This dose of tobramycin was choosen because it causes a relatively small change in *P. aeruginosa* (∼1 log_10_) observed in clinical studies, including one study with a single acute exposure to tobramycin (19–21). Figure 1 reveals that tobramycin reduced the percent of live bacteria from 94.0% in control to 88.9% in the presence of tobramycin. The percent of dead *P. aeruginosa* increased from 6.0% in control to 11.1% in tobramycin exposed bacteria. This change in live/dead *P. aeruginosa* is similar to the effect of tobramycin on *P. aeruginosa* in clinical samples obtained from pwCF (19–21). Tobramycin also reduced the growth of *P. aeruginosa* in an *in vitro* assay (Figure 1).

**Figure 1.**
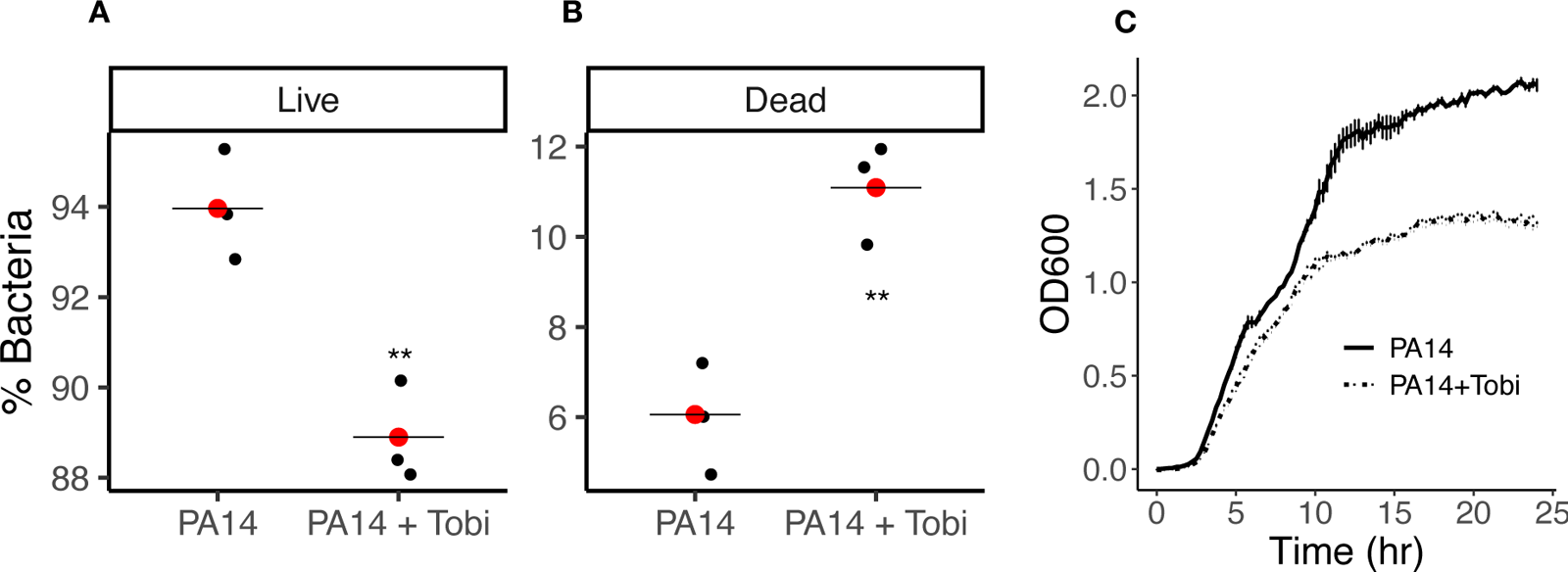
(**A,B**) The LIVE/DEAD *Bac*Light Viability Kit™ revealed that tobramycin (1 μg/ml) reduced the viability of *P. aeruginosa*. A t test was used to determine significance. The means are presented as red dots. Each black dot represents an experiment with a unique culture of PA14 exposed to vehicle or tobramycin on different days. (**C**) Tobramycin (1 μg/mL; PA14 + Tobi) reduced growth of PA14 as determined by measuring OD600. Growth curve of PA14 in the presence of vehicle or tobramycin. Lines represent the averages from three biological replicates conducted of different days, and error bars (too small to see) indicate SEM. The growth curves first significantly diverge at 1.15 hours (P<0.05, t test) and remain significantly different thereafter. **P<0.01

### Isolation and characterization of OMVs

OMVs secreted by *P. aeruginosa* were isolated using the OptiPrep™-Density Gradient. To determine the fraction of the density gradient containing OMVs the column was divided into 9 equal fractions (0.5 mL each), whereupon the number of OMVs in each fraction was assessed by Nanosight Tracking Analysis (NTA). OMVs secreted by *P. aeruginosa* exposed to tobramycin (Tobi-OMVs) were most frequent in fractions 2 and 3 and OMVs secreted by *P. aeruginosa* exposed to vehicle (V-OMVs) were also most frequent in fractions 2-3 (Figure 2). Thus, fractions 2-3 were combined in each treatment group for all subsequent studies. Tobramycin increased the number of secreted OMVs by 38%, compared to V-OMV (Figure 2B). The increase in OMV numbers in the tobramycin exposed *P. aeruginosa* is likely due in part to hypervesiculation and/or due to membrane fragments released from dead bacteria (48), an effect that is also likely to occur in *in vivo* in pwCF treated with tobramycin.

**Figure 2.**
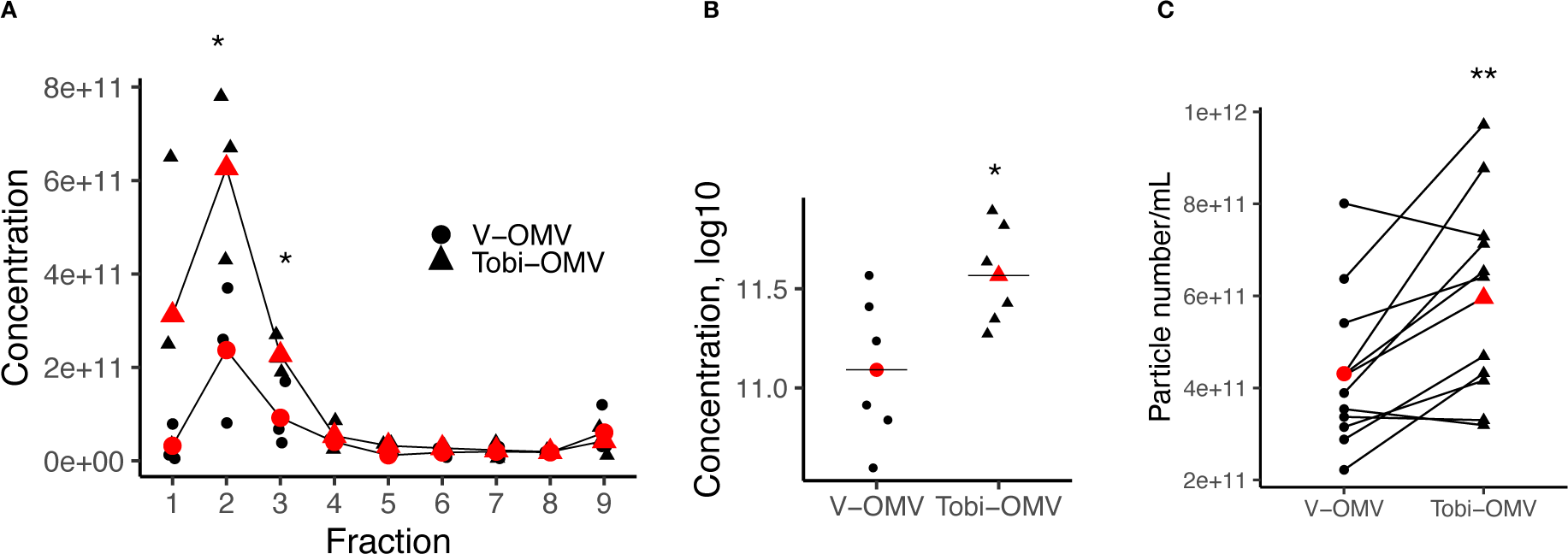
Nanosight particle tracking analysis (NTA) of OMVs isolated using the OptiPrep™-Density Gradient. (A) NTA of OMVs detected in each OptiPrep™-fraction from six separate experiments conducted on different days. Vehicle exposed (V-OMV) *P. aeruginosa*, and tobramycin exposed *P. aeruginosa* (Tobi-OMV) are shown. The red dots indicate the means. The number of Tobi-OMVs was significantly above V-OMVs in fractions 2 and 3, thus, OMVs were pooled from fractions 2-3 from V-OMV and fractions 2-3 were pooled from Tobi-OMV. N=6 experiments conducted on separated days with different grow ups of *P. aeruginosa*. (B) Total counts of OMVs in fractions 2 and 3 combined from the same experiments depicted in A (N=6). (C). Number of V-OMVs and Tobi-OMVs in 12 independent experiments conducted on different days where fractions 2 and 3 were combined for reported experiments. Each bacterial preparation was separated into two aliquotes, one was treated with vehicle and the other tobramycin, and then OMVs were isolated as described in Methods. The lines connect experiments on the same day. *P<0.05, **P<0.01.

OMVs were also analyzed by negative staining electron microscopy (Figure 3). OMVs were observed in V-OMV samples and Tobi-OMV samples, but not in process control samples (media not exposed to *P. aeruginosa* and processed through the OMV isolation procedure to account for any possible effects of the isolation process on HBEC)(Figure 3). V-OMVs were 28.8 ± 6.5 nm (SD) in diameter (n=322, range 18-70 nm). Tobi-OMVs were 27.2 ± 8.4 nm (SD) (n=326, range 20-80 nm). Tobramycin had no effect on the size of OMVs (unpaired t test). Both values are similar to other reports in the literature on the diameter of OMVs secreted by *P. aeruginosa* (49, 50).

**Figure 3.**
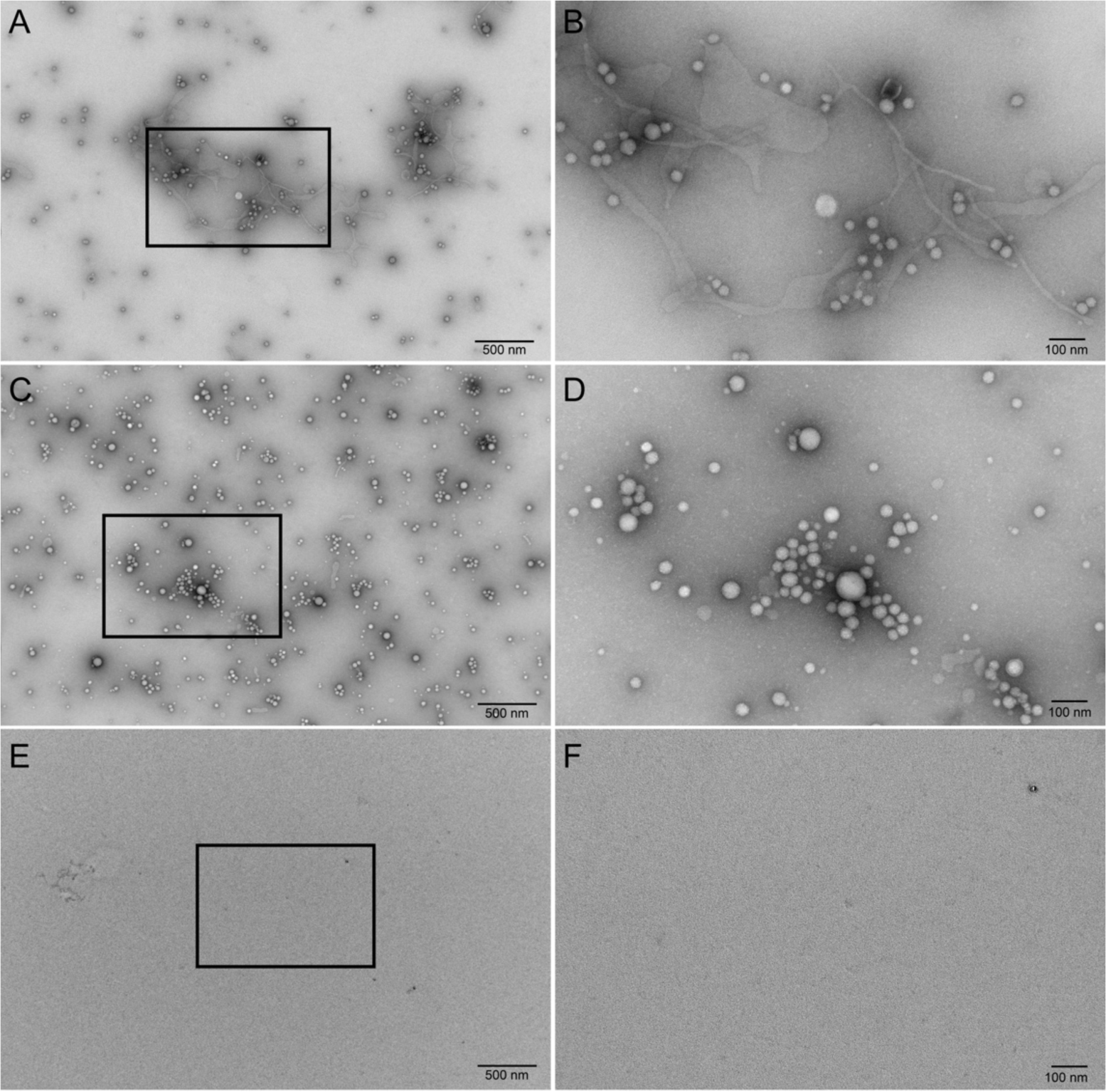
Representative transmission electron microscopy images of negatively stained OMV preparations and process control. The left-hand column shows images acquired at an original magnification of 10,000X, and the right-hand column shows the boxed areas from the left-hand column images acquired at an original magnification of 30,000X. Scale bars represent 500 nm and 100 nm for the low and high magnifications, respectively. (A,B) Images of V-OMVs: (A) low magnification and (B) high magnification. (C,D) Images of Tobi-OMVs: (C) low magnification and (D) high magnification. (E,F) Images showing the lack of identifiable vesicles in process control (media not exposed to PA14 and run through the OMV isolation procedure). (E) low magnification and (F) high magnification. Experiments repeated twice on separate days. The electron microscopist was blinded to the treatment.

### V-OMVs elicited a more robust immune reponse than Tobi-OMVs

To examine the effect of OMVs on the host immune response, CF-HBECs from CF donors (homozygous for Phe508del, the most common mutation in CFTR) were grown in air-liquid interface (ALI) culture (31, 38) and exposed to vehicle, V-OMVs or Tobi-OMVs for 6 hours, whereupon the secretion of 48 cytokines was measured by ELISA. IL-8 (Figure 4A), a neutrophil chemoattractant, and IP-10 (Figure 4B), a chemoattractant for monocytes, macrophages and other immune cells, were the only cytokines significantly increased by V-OMVs and Tobi-OMVs compared to vehicle control and significantly different between V-OMV and Tobi-OMV (Figure 4). Both IL-8 and IP-10 secretion were less in CF-HBEC exposed to Tobi-OMVs compared to V-OMVs (Figure 4).

**Figure 4.**
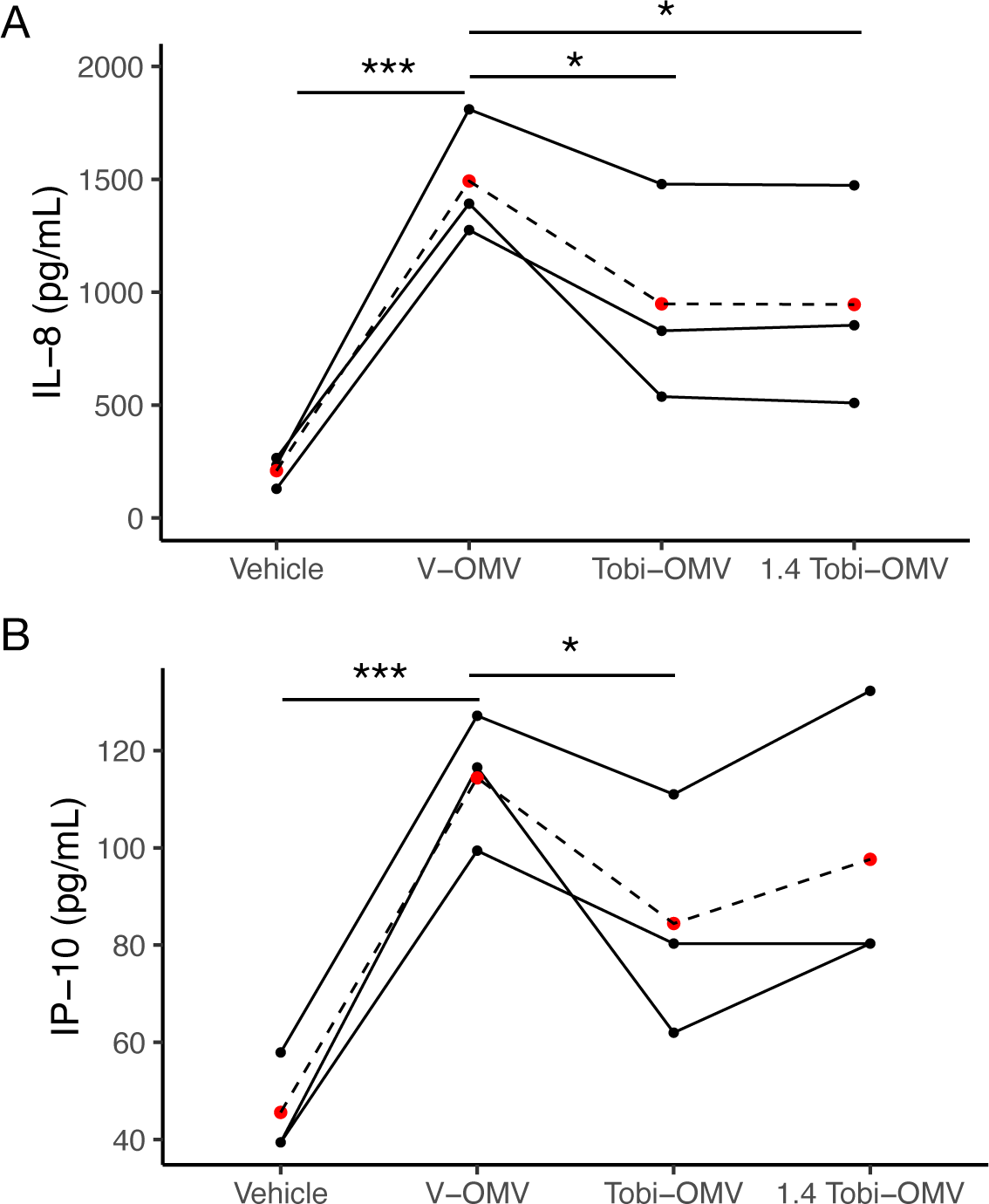
CF-HBECs from three donors were exposed to either vehicle, the same number of V-OMVs, or 40% more Tobi-OMVs, to represent the increased secretion of OMVs induced by tobramycin in the *in vitro* experiments. The concentration of OMVs used was in the range of bacterial vesicles measured in biological fluids, including BALF, *in vivo* [reviewed in (28, 29)]. After a six-hour exposure, the basolateral medium was collected and cytokines were interogated by 41-plex ELISA. Cytokines other than IL-8 and IP-10 were not significantly different between V-OMV and Tobi-OMV. Lines connect experiments conducted with CF-HBECs from the same donor on the same day. (**A**) V-OMVs increased IL-8 secretion compared to vehicle. Tobi-OMVs induced less IL-8 secretion compared to V-OMVs. 1.4 times as many Tobi-OMVs (2.1 x 10^10^ for a 12-mm filter) had a similar effect on IL-8 secretion compared to Tobi-OMVs (1.5 x 10^10^ for a 12-mm filter). \**P* < 0.05; \*\*\**P* < 0.001. **(B)** V-OMVs increased IP-10 secretion compared to vehicle. Tobi-OMVs induced less IP-10 secretion compard to V-OMVs. Lines connect experiments conducted with CF-HBECs from the same donor. Linear mixed-effects models with CF-HBEC donor as a random effect were used to calculate *P* values. \**P* < 0.05. \*\*\**P* < 0.001

### Tobramycin increases the abundance of tRNA-fMet halves in OMVs

To test the hypothesis that tobramycin increased the abundance of anti-inflammatory sRNAs in OMVs, a small RNA-sequencing analysis was performed to compare the sRNA content in V-OMVs and Tobi-OMVs. We identified 1064 unique sequences that were differentially enriched in Tobi-OMVs compared to V-OMVs. Thus, studies were focused on differentially induced sRNAs between V-OMV and Tobi-OMV exposed CF-HBEC for additional study (Figure 5A and Table 1). We chose two 35-nt long sRNAs (#5 and #7 in Table 1) that were fragments of two initiator tRNAs (tRNA-fMet1 and tRNA-fMet2 located at PA14_62790 and PA14_52320, respectively) in PA14 for further analysis because they were predicted by miRanda to suppress IL-8 secretion by CF-HBECs. The sequence reads of the tRNA-fMet halves in Tobi-OMVs mapped to these two loci had similar length distributions (Figure 5B and 5C; 80% of reads were 35-nt long). Both have low minimum free energy, suggesting stable secondary structures. The two 35-nt long tRNA-fMet fragments are 5′ halves of tRNA-fMet halves (hereafter called tRNA-fMet halves), which are products of cleavage in the anticodon loop (Figure 5D). Importantly, the two tRNA-fMet halves have high sequence similarity with only one nucleotide difference, suggesting similar sequence-based targeting functions. The high sequence similarity allowed us to design quantitative PCR (qPCR) primers to quantify both tRNA-fMet halves simultaneously. By qPCR, we confirmed the RNA-seq data that tobramycin increased the amout of tRNA-fMet halves in OMVs secreted by PA14 (Figure 5E). In addition, tobramycin also increased the amout of tRNA-fMet halves in OMVs secreted by four CF clinical isolates of *P. aeruginosa* (Figure 5E), indicating a bacterial strain-independent phenotype. Moreover, we reanalyzed our published small RNA-sequencing experiment (36), in which we reported transfer of a PA14 sRNAs (24 nt) into HBECs after V-OMVs exposure, and we identified both tRNA-fMet halves in OMV-exposed HBEC but not in vehicle control HBEC (36). Taken together, these observations demonstrate that the two 35 nt tRNA-fMet halves are differentially induced sRNAs in Tobi-OMVs, have high sequence similarity, and are delivered to airway epithelial cells by OMVs; thus, they are good candidates for further investigation into their possible role in suppressing IL-8 secretion.

**Figure 5.**
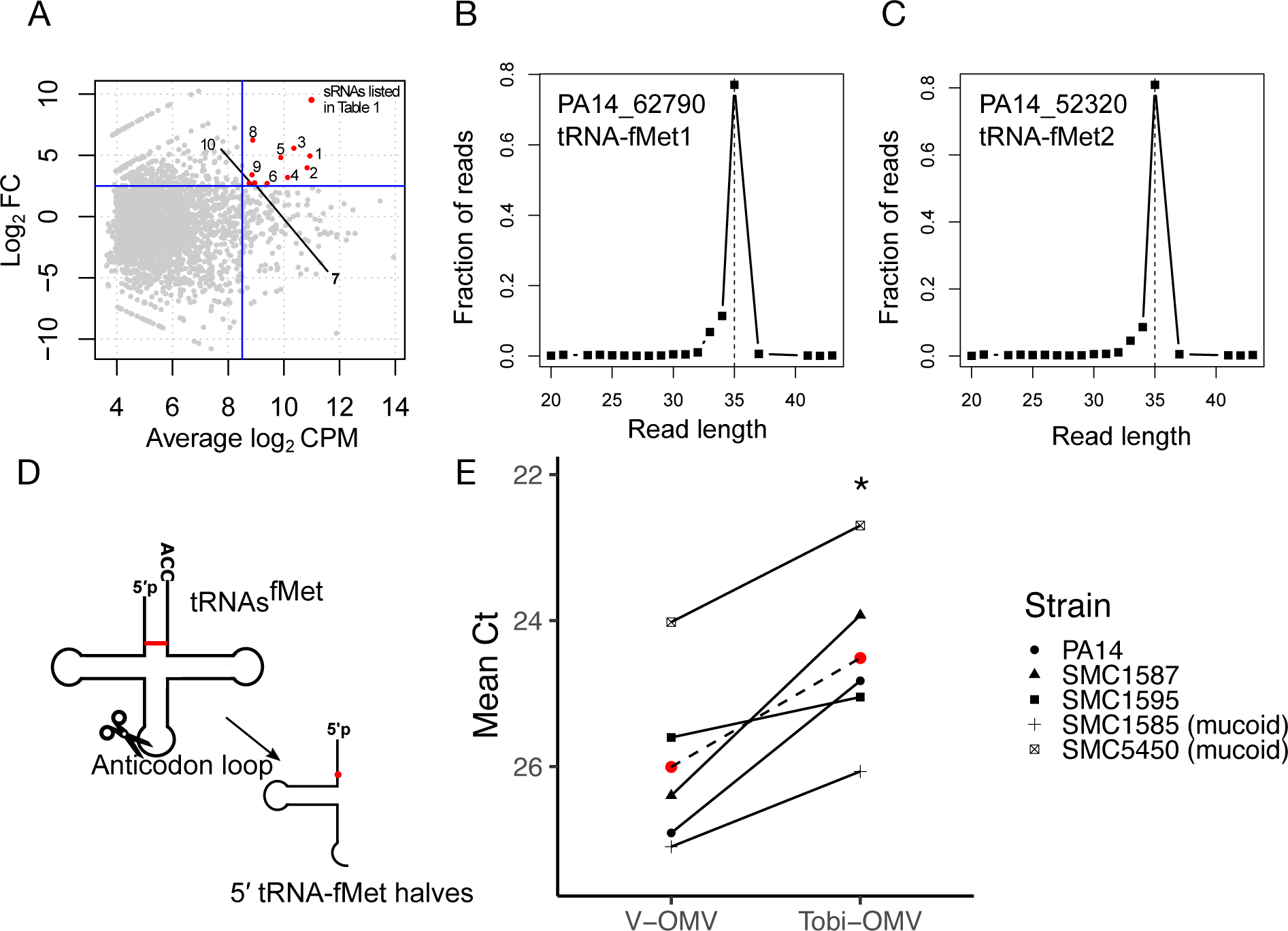
Tobramycin increases the abundance of tRNA-fMet halves in OMVs. The input of small RNAs into the sequencing process was similar in each group (see Methods). (**A**) M versus A plot (MA plot; M is the difference between the log intensity values and A is the average of the log intensity values), comparing the small RNA expression profile in V-OMVs and Tobi-OMVs (N = 3 for each group). Each dot represents a unique sequence read. The most abundant and most induced sRNAs in OMVs by tobramycin are highlighted in red and listed in Table 1. (**B and C**) Length distribution of tRNA-fMet halves secreted in Tobi-OMVs mapped to gene lous *PA14_62790* (**B**) and *PA14_52320* (**C**). (**D**) Secondary cloverleaf structure of tRNA-fMet and cleavage site in the anticodon loop to generate tRNA-fMet halves. The red line indicates the only different pair of nucleotides between the two 5′ tRNA-fMet halves, and the red dot represents the only nucleotide difference between the two 5′ tRNA-fMet halves. (**E**) qPCR for 5′ tRNA-fMet halves in V-OMVs and Tobi-OMVs purified from PA14 and four CF clinical isolates (N = 5 strains), including two mucoid and two non-mucoid CF clinical isolates. The qPCR primers and probe were designed to detect both 5′ tRNA-fMet halves. The red dots indicate the mean. A t-test was used to establish significance. * *P* < 0.05.

**Table 1.** Top 10 most abundant and most differentially induced sRNAs in Tobi-OMVs compared to V-OMVs.

| # | PA14 locus | Gene product of the locus | Log2FC | Average Log2CPM | length | Minimum free energy (kcal/mol)<sup>a</sup> |
| --- | --- | --- | --- | --- | --- | --- |
| 1 | Multiple | tRNA-Asp | 4.95 | 10.93 | 23 (20, 23)<sup>b</sup> | -0.2 |
| 2 | Multiple | 16S rRNA | 3.98 | 10.83 | 33 (30-39)<sup>b</sup> | -6.9 |
| 3 | Multiple | 23S rRNA | 5.57 | 10.35 | 44 | -14.6 |
| 4 | Multiple | tRNA-Ala | 3.19 | 10.13 | 34 | -8 |
| 5 | 62790 | tRNA-fMet1 | 4.82 | 9.88 | 35 | -7.5 |
| 6 | 28740 | tRNA-Pro | 2.70 | 9.39 | 36 | -9.7 |
| 7 | 52320 | tRNA-fMet2 | 2.74 | 8.95 | 35 | -9.3 |
| 8 | 61760 | tRNA-Gln | 6.23 | 8.88 | 20 | -0.2 |
| 9 | 30720 | tRNA-Cys | 3.41 | 8.85 | 40 | -4.6 |
| 10 | multiple | 5S rRNA | 2.72 | 8.75 | 45 | -3.8 |

<sup>a</sup>The minimum free energy for each sRNA was predicted using the RNAfold web server (78).
<sup>b</sup>Multiple reads of different lengths were mapped to the same locus, and the most abundant reads are listed in Table 1.

### tRNA-fMet halves reduce IL-8 secretion by CF-HBEC

To determine if tRNA-fMet halves reduce IL-8 secretion, we transformed PA14 with an arabinose-inducible vector expressing the tRNA-fMet1 half (tRNA1-OMVs) or an empty vector control RNA (EV-OMV). Q-PCR experiments confirmed that the expression of tRNA-fMet1 halves in tRNA1-OMVs secreted by PA14 was significantly induced (2.73 fold) compared to EV-OMVs secreted by control exposed PA14 (Supplemental Figure 1). CF-HBECs were exposed to EV-OMVs or tRNA1-OMVs for 6 hours, and the secretion of IL-8 was measured by ELISA. As predicted by miRanda, tRNA1-OMVs induced less IL-8 secretion compared to the same amount of EV-OMVs (Figure 6A). To provide additional support for the conclusion that tRNA-fMet halves reduce IL-8 secretion by CF-HBECs, we designed an inhibitor, an RNA oligonucleotide with a complementary sequence to both tRNA-fMet halves. CF-HBECs were transfected with a control RNA oligonucleotide, not predicted to target any genes in *P. aeruginosa* (ThermoFisher Scientific) or an antisense RNA oligo inhibitor of the tRNA-fMet halves followed by exposure to V-OMVs or 1.4X Tobi-OMVs. As predicted, the antisense RNA oligonucleotide inhibitor reduced the ability of Tobi-OMVs to suppress IL-8 secretion compared to V-OMVs, whereas the control RNA oligonucleotide had no effect on the OMV response (Figure 6B). Thus, these experiments, taken together, demonstrate that the tRNA-fMet halves in OMVs reduce OMV-stimulated IL-8 secretion.

**Figure 6.**
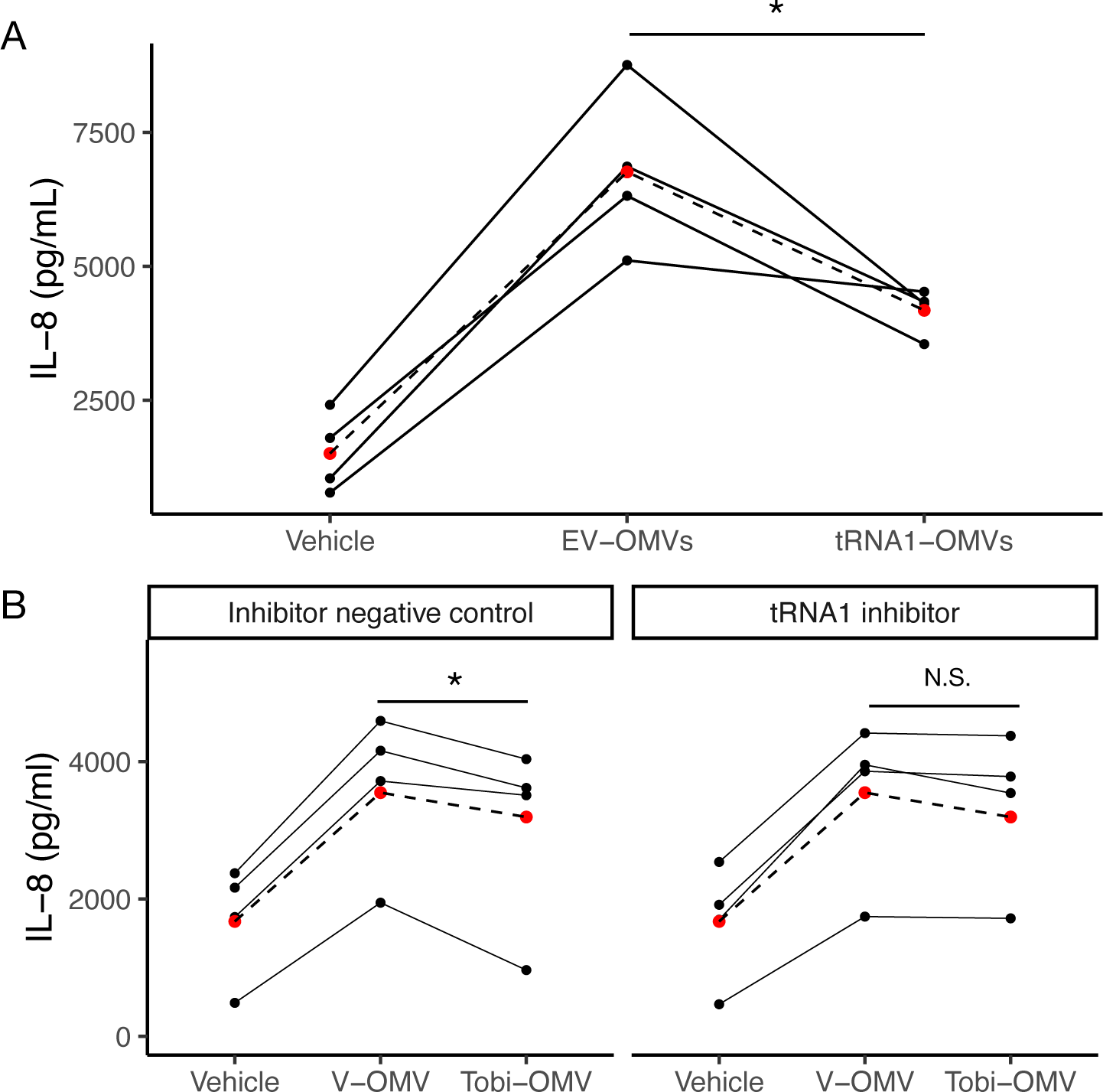
tRNA-fMet halves reduce IL-8 secretion *in vitro.* **(A)** Empty vector OMVs (EV-OMV) increased IL-8 secretion by CF-HBECs (N = 4 donors). OMVs overexpressing tRNA-fMet1 half (tRNA1-OMVs) reduced IL-8 secretion compared to EV-OMV. **(B)** The 1.4X Tobi-OMV effect of reducing IL-8 secretion was abolished by transfection of an antisense RNA oligo inhibitor (tRNA1 inhibitor) in CF-HBEC that anneals to both tRNA-fMet halves but not by transfection of inhibitor negative control (n = 4 donors). Lines in panels A and B connect data points using the cells from the same donor on the same day. A linear mixed-effects model with CF-HBEC donor as a random effect was used to calculate *P* values for data in A and B. *P<0.05.

### Identification of tRNA-fMet1 half targets in HBEC

To identify potential gene targets of the tRNA-fMet halfs in CF-HBEC, a miRanda microRNA target scan was performed (42). miRanda is an algorithm designed for RNA-RNA binding predictions considering sequence complementarity and binding free energy. Given the high sequence similarity between the two sRNAs, we used the sequence of tRNA-fMet1 half to scan the whole human transcriptome and adjusted the prediction for the gene expression profile of CF-HBEC and generated a list of 1518 predicted targets, accounting for 8.4% of human coding genes. From this list of 1518 targets Ingenuity Pathway Analysis (IPA) identified several pro-inflammatory pathways in epithelial cells that are predicted to be down-regulated by the tRNA-fMet1 half and reduce IL-8 secretion (Table 2).

**Table 2.**
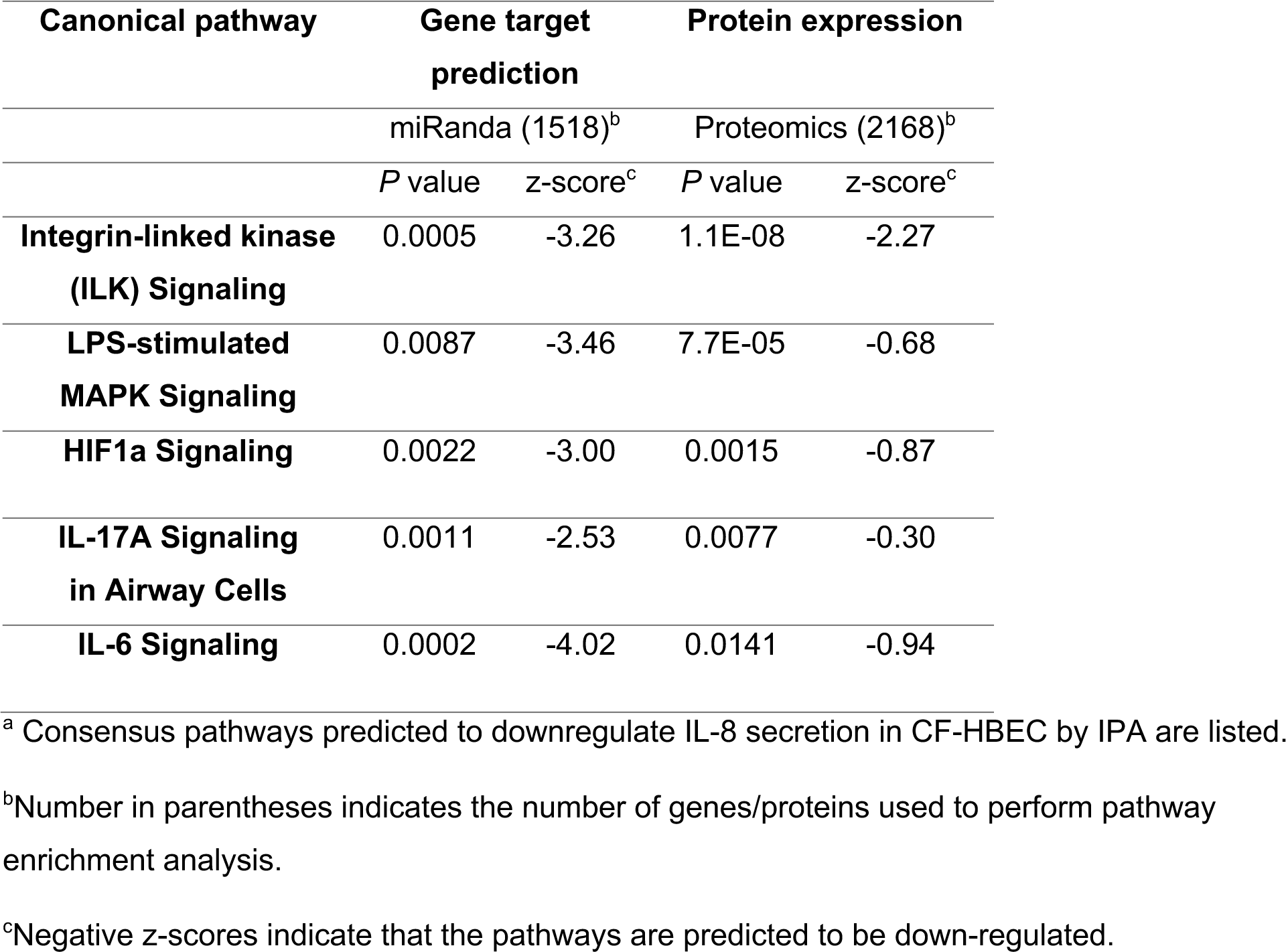
Significantly enriched signaling pathways predicted to regulate IL-8, identified using miRanda and IPA analysis. Proteomic analysis of CF-HBEC confirmed that proteins in the canonical pathways were down-regulated^a^.

To identify proteins whose abundances were changed by tRNA-fMet halves and are in the predicted pathways identified by IPA, CF-HBECs were exposed to EV-OMVs or tRNA1-OMVs for 6 hours before being subjected to proteomic analysis. 8343 proteins were identified, and the top 20% of differentially expressed proteins were selected for further analysis, yielding 943 significantly down-regulated proteins (Figure 7A). Several of the pathways identifed by IPA analysis of the differentially expressed genes are pro-inflammatory and induce IL-8 secretion (Table 3)(51). Specifically, IPA analysis identified seven proteins including MAPK10, IKBKG, and EP300 that were decreased in the proteomics experiments, suggesting tRNA-fMet halves targeting of a pro-inflammatory network, resulting in the reduction of IL-8 secretion (Figure 7B). In summary, these experiments demonstrate that tRNA-fMet halves transferred from OMVs to CF-HBECs are predicted by miRanda to target several genes that mediate the Tobi-OMV induced reduction in IL-8 secretion compared to V-OMV. Moreover, the proteomic analysis confirmed that the tRNA-fMet half reduced MAPK10, IKBKG, and EP300, signaling molecules that play a key role in increasing IL-8 secretion (Figure 7).

**Figure 7.**
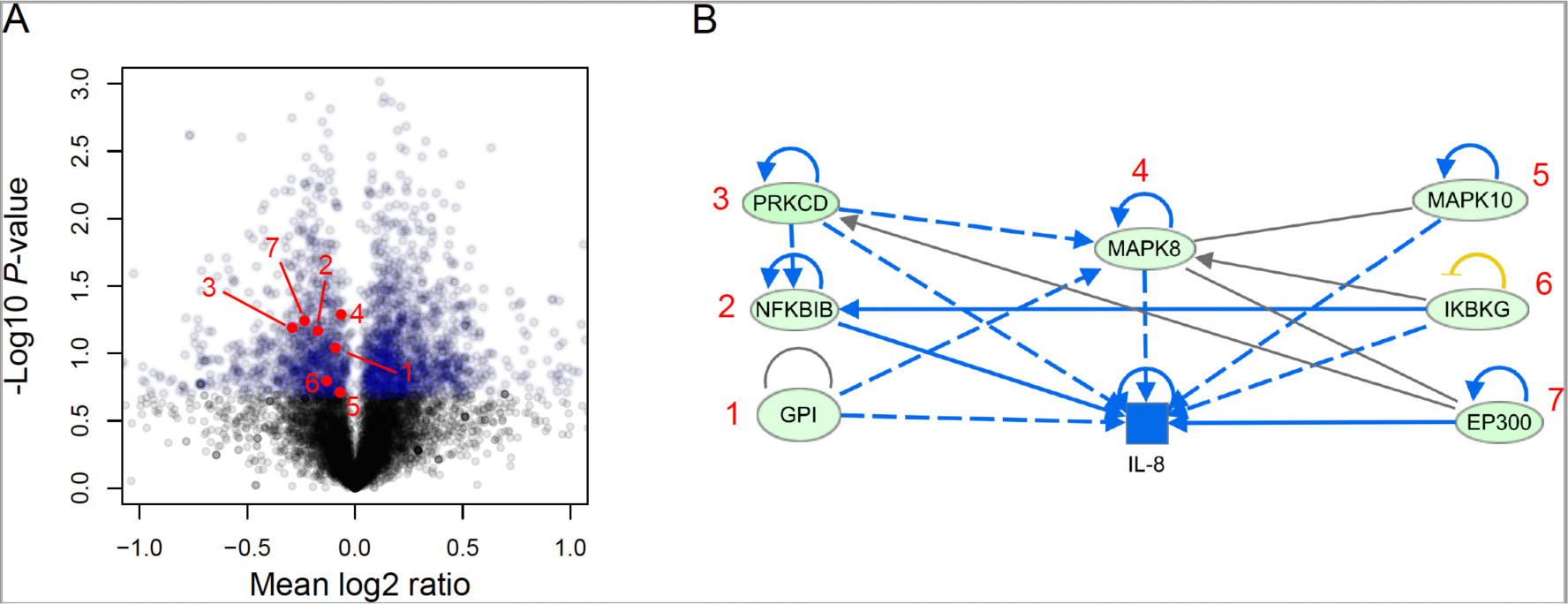
tRNA-fMet halves alter protein expression in CF-HBEC. (**A**) Volcano plot of proteomic analysis of polarized CF-HBECs (n = 3 donors) exposed to tRNA-fMet1 half (tRNA1-OMVs) compared to cells treated with EV-OMVs. The top 20% differentially expressed proteins are colored in blue (FDR P<0.05 and >2-fold increase in abundance). Red dots with numbers represent down-regulated proteins corresponding to proteins numbered in B. (**B**) Ingenuity Pathway analysis (IPA) identified a down-regulated pro-inflammatory network in five consensus pathways (Table 3), predicting decreased IL-8 expression. The green circles identify proteins whose abundance was reduced based on proteomic analysis. Blue shading indicates predicted inhibition.

### Tobramycin Reduces the Pro-inflammatory Effect of OMVs in Mouse Lungs

Studies were also conducted in mice to further support the conclusion that tobramycin reduces the pro-inflammatory effect of OMVs by increasing the tRNA-fMet halves content. Mice were exposed to EV-OMVs or tRNA1-OMVs for 6 hours, and BALF was harvested for analysis. The concentration of KC, a murine functional homolog of IL-8 (Figure 8A), and neutrophil content (Figure 8B) were significantly reduced in BALF obtained from mice exposed to tRNA1-OMVs compared to EV-OMVs. Thus, the increse in the abundance of tRNA-fMet halves in Tobi-OMVs reduced the pro-inflammatory response in a mouse model of infection and inflammation compared to V-OMVs. Studies in which we attempted to delete the tRNA-fMet halves in *P. aeruginosa* was lethal to the bacteria, thus, we were unable to conduct tRNA-fMet halves deletion experiments in *P. aeruginosa*.

**Figure 8.**
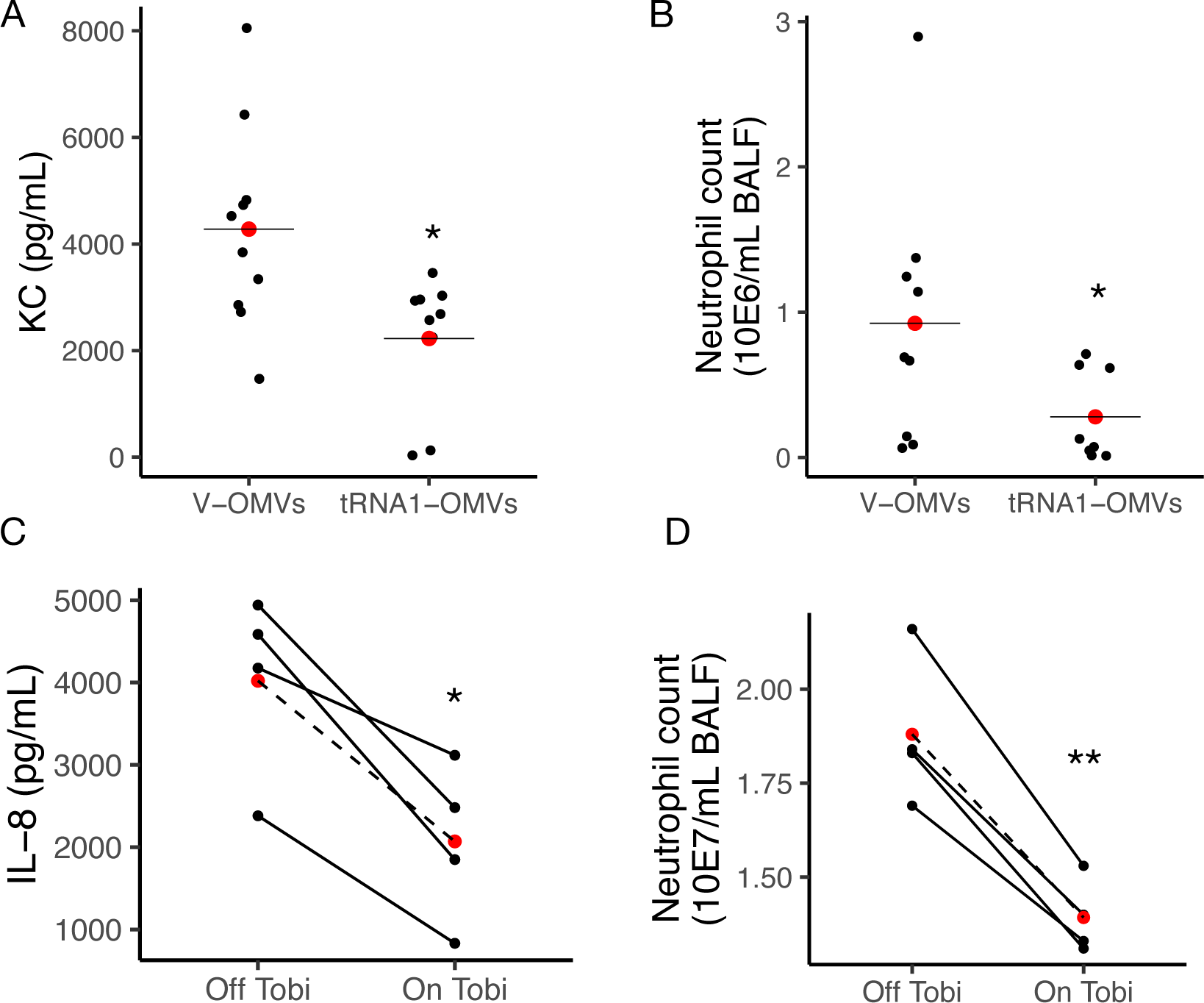
tRNA-fMet halves reduce IL-8 secretion and neutrophil recruitment *in vivo.* BALF from male and female mice exposed to EV-OMVs or tRNA1-fMet half OMVs was collected to measure KC concentration (A) and neutrophils (B). 9 to 10 mice were used per group, and the Wilcoxon rank-sum test was used to test significance. (C and D) BALF samples collected from four pwCF during the 4-week administration of inhaled tobramycin (On Tobi) or not (Off Tobi). Lines connect data points from the same pwCF. Linear mixed-effect models were used to account for donor-to-donor variability and the number of days between collection dates for each sample pair (Supplemental Table 1). Red dots indicate the mean.\**P* < 0.05; \*\**P* < 0.001. Results were independent of the order of sample collection. The Dartmouth College IRB will not approve consecutive, prospective bronchoscopies in pwCF for research purposes; thus samples were not collected in consecutive months. The samples collected and reported herein represent clinically justified samples of BALF to be used for research approved by the Dartmouth IRB.

### Inhaled tobramycin has an anti-inflammatory effect in *P. aeruginosa*-infected human CF lungs

To determine if tobramycin exposure is associated with less inflammation and neutrophil burden in pwCF, compared to the period off tobramycin, we performed a retrospective analysis to assess whether the administration of inhaled tobramycin changes the inflammatory status in pwCF. Bronchoalveolar lavage fluid (BALF) samples were collected from four pwCF chronically infected with *P. aeruginosa* during the month of inhaled tobramycin (On Tobi) and the month off tobramycin (Off Tobi). In BALF obtained On Tobi, IL-8 levels were reduced by 48.5% (Figure 8C), and the number of neutrophils were decreased by 25.9% (Figure 8D) compared to BALF collected Off Tobi. This clinical correlation is consistent with the *in vitro* CF-HBEC and *in vivo* mouse experiments.

## Discussion

The goal of this study was to determine how tobramycin might improve clinical outcomes in pwCF without significantly or modestly reducing the abundance of *P. aeruginosa* in the lungs. Our data reveal that tobramycin increases the concentration of 35 nt tRNA-fMet halves in OMVs secreted by *P. aeruginosa*, that the OMVs deliver tRNA-fMet halves to CF-HBECs, and that the increased delivery of tRNA-fMet halves to CF-HBECs suppresses OMV-mediated IL-8 secretion by down regulating MAPK10, IKBKG, and EP300 protein abundance, key proteins involved in the IL-8 secretory pathway. Both *in vitro* experiments on primary CF-HBEC from four donors and *in vivo* experiments in mice are consistent with this conclusion. Our retrospective analysis of pwCF on and off tobramycin is consistent with our data in mice that tobramycin reduces IL-8 and the neutrophil content in BALF, although it should be noted that it is not possible to definatively state that the anti-inflammatory effect of tobramycin in pwCF is mediated by an increase in the tRNA-fMet half in OMVs. The reduction in the neutrophil content in BALF is predicted to mitigate damage in the CF lungs since CF neutrophils are a source of significant lung damage in pwCF (5, 6). Additional studies, beyond the scope of the present study, are required to determine if the magnitude of the change in the tRNA-fMet secreted in Tobi-OMVs is sufficient to migitate lung damage in mice casued by *P. aeruginosa* infection.

In a previous study we demonstrated that OMVs secreted by *P. aeruginosa* induced a pro-inflammatory response (increased IL-8) in HBEC and mouse lung (increased KC) that was mitigated by a 24 nt siRNA fragment of sRNA52320 that was abundant inside OMVs, resistant to RNases and transferred into HBEC (36). Taken together with the present study, we have shown that multiple sRNAs are present in OMVs, including the 35 nt tRNA-Met halves identified in the present study and the 24 nt sRNA sRNA52320 that are capable of selectively down-regulating IL-8/KC levels in mice BALF and IL-8 and IP-10 secreted by CF-HBEC.

The most parsimonious interpretation of our past and current data is that pathogen-associated molecular patterns (PAMPs) on the outside of OMVs secreted by *P. aeruginosa* stimulate the host innate immune response by activating Toll-like receptors (TLR). Whereas the presence of the tRNA-Met half fragments in OMVs mitigates the pro-inflammatory effect of OMV PAMPs. In addition, tobramycin, by upregulating the tRNA-Met half content of OMVs decreases the IL-8 and neutrophil levels in the lungs of mice. Given that the classic TLR-mediated innate immune response to LPS involves the up-regulation of multiple pro-inflammatory cytokines, it is surprising that only IL-8/KC and IP-10 were repressed by the tRNA-Met halves in the present study. One explanation for this could be that the tRNA-Met halves targets mRNA in the TLR2/4-induced innate immune response pathway, while cytokine secretion mediated by other receptors and pathways remains unaffected. For example, knockout of TLR4 and TLR2 blocks the OMV-induced secretion of KC more than any other cytokine (52). Inhibition of TLR2 and TLR4 decreases *Mycobacterium bovis*-induced ERK1/2 activation and subsequent IL-8 secretion in human epithelial cells (53). The inhibition of IL-8 and KC secretion by the tRNA-Met halves that we observed in this study is consistent with the hypothesis that the tRNA-Met halves primarily attenuate TLR4 signaling. Thus, the data in this report suggest a new explanation for how long-term tobramycin administration improves lung function in pwCF in addition to the bactericidal effect of tobramycin. Since aminoglycoside antibiotics are also bactericidal to other bacteria in addition to *P. aeruginosa*, the effect of tobramycin in pwCF could also be due to a decrease in the number and/or virulence of other bacteria in pwCF (21).

tRNA-derived fragments are a novel class of regulatory sRNAs in prokaryotes and eukaryotes (27, 54). tRNAs are the most abundant RNA species by the number of molecules, and fragments with different lengths have been reported in three domains of life. miRNA-sized (∼24-nt long) tRNA fragments in mammalian cells have garnered attention as they have been found to associate with Argonaute (AGO) proteins to mediate gene silencing by base-pairing with target mRNAs (55–57). Recently, a report showed that in *Bradyrhizobium japonicum*, a 21-nt tRNA fragment utilizes host plant AGO1 to regulate host gene expression, a cross-kingdom symbiotic relationship between bacteria and plants (58). tRNA halves have been shown to have both positive and negative effects on translation by regulating the formation of ribosomes and the translation initiation complex (59–62). For example, *Helicobacter pylori* secretes sR-2509025, a 31-nt 5′ tRNA-fMet fragment, in OMVs that fuse with human gastric adenocarcinoma cells and diminishes LPS-induced IL-8 secretion (63). Additional studies, beyond the scope of the present study are required to determine if the tRNA-fMet halves secreted by *P. aeruginosa* in OMVs suppress inflammation in CF-HBEC by an AGO-dependent mechanism.

Numerous reports have demonstrated that NF-κB and MAPK signaling pathways induce IL-8 secretion (64–66). IKBKG, also known as NF-κB essential modulator (NEMO), is critical for NF-κB pathway activation. EP300, also called P300, is a transcription co-factor required for NF-κB-dependent IL-8 induction (67, 68). Moreover, a study demonstrated that DNA damage lead to NF-κB activation followed by MAPK10-mediated IL-8 secretion (69). Indeed, the elevated DNA damage response correlates with the non-resolving neutrophilic inflammation in the CF airways (70, 71). Hence, our findings reveal that tRNA-fMet halves decrease IKBKG, EP300, and MAPK10 protein expression thereby reduce IL-8 secretion and neutrophil levels remain consistent with the literature on the NF-κB pathway. Here, we demonstrate that tRNA-fMet halves target a pro-inflammatory network involving the MAPK and NF-κB signaling pathways, which are intrinsically over-activated in CF (3, 4), highlighting the importance of this network in pulmonary inflammation.

Our study has several advantages, including: i) The use of primary CF-HBEC from multiple donors is more representative of the human population, which manifests considerable heterogeneity, than immortalized or tumor cell lines isolated from a single donor or the use of non-epithelial cell types (72), ii) The use of four CF clinical isolates of *P. aeruginosa* (mucoid and non-mucoid), demonstrating that the ability of tobramycin to increase the abundance of the tRNA-fMet halves in OMVs is not strain dependent, iii) Demonstration that increased expression of the tRNA-fMet halves in *P. aeruginosa* decreased the ability of OMVs to increase IL-8 and neutrophils in mouse BALF compared to EV-OMV, and iv) The observation interrogating BALF samples obtained from pwCF are also consistent with our CF-HBEC and mouse studies. Thus, several lines of evidence using a variety of approaches are consistent with our conclusion that tobramycin increases the concentration of tRNA-fMet halves secreted by *P. aeruginosa* in OMVs, and that the increase in tRNA-fMet halves abundance mitigates OMV-induced inflammation. There are a few limitations of our study. First, we performed a retrospective analysis of BALF samples collected from pwCF on and off inhaled tobramycin; however, we could not collect BALF in consecutive months on and off tobramycin in the same individuals because of the invasive nature of the technique and IRB restrictions on research bronchoscopies at Dartmouth Health. Nevertheless, after adjusting for the number of days between collection dates and the order of sample collections for each sample pair (range from 175 to 791 days (Supplemental Table 1), tobramycin-on BALF had significantly lower IL-8 concentration and fewer neutrophil counts than tobramycin-off BALF. Importantly, a similar observation was made for IL-8 in CF sputum samples collected in consecutive months from pwCF on and off tobramycin (73). We acknowledge that the studies on BALF isolated from pwCF does not prove that tobramycin increased the secretion of tRNA-fMet in OMVs in pwCF and that a change in the tRNA-fMet was responsible for the *in vivo* effects of tobramycin in pwCF. Because studies have shown that IL-8 concentration in sputum is inversely correlated with pulmonary function (74, 75), we speculate that inhaled tobramycin has an anti-inflammatory effect in *P. aeruginosa*-infected CF lungs, resulting in a reduction in neutrophils and improved lung function. Additional studies are needed to evaluate the effect of the tRNA-fMet halves on lung function. Second, since there are many other differences in the sRNA content and the virulence factor content of Tobi-OMVs compared to V-OMVs (30) we cannot rule out the possibility that other factors may contribute to the difference in the immune response of CF-HBECs and mouse lungs to Tobi-OMVs versus V-OMVs. Nevertheless, since an inhibitor of tRNA-fMet halves transfected into CF-HBECs blocked the Tobi-OMV mediated reduction in IL-8 secretion compared to V-OMV, we conclude that tRNA-fMet halves play an important role in suppressing the OMV-induced increase in IL-8 and neutrophil levels. Third, tRNAs are known to have post-transcriptional modifications, which affect RNA structure and RNA-protein interaction (76). Whether or not modifications on tRNA-fMet halves modulate the anti-inflammatory effect will require additional studies. Finally, in our *in vivo* studies on mouse and pwCF we consider it likley that the tRNA-Met halves may also decrease IL-8 secretion by immune cells in the lungs as well as CF-HBEC. More studies are required to examine the effect of the tRNA-Met halves on immune cells.

Highly effective CFTR modulator drugs have significantly improved outcomes in pwCF; however, in a few recent studies, they have been shown to have either no effect or a modest effect on the *P. aeruginosa* burden in the CF lungs and on the hyperinflammatory state (7, 19–21, 77). Thus, new approaches are needed to reduce the bacterial load and excessive inflammation in the lungs of pwCF. We propose that tRNA-fMet halves or similar miRNA-like molecules may be utilized as a therapeutic strategy to reduce IL-8 and neutrophil content in the lungs of pwCF, resulting in reduced lung damage and, therefore, improved lung function.

## Acknowledgments

This work was supported by the Cystic Fibrosis Foundation (STANTO19G0, STANTO20PO, STANTO19R0, and HOGAN19G0), the NIH (P30-DK117469, R01HL151385, P20-GM113132, S10OD016262), the Dartmouth Cancer Center Core Grants (5P30 CA023108-41, P30CA023108) and the Flatley Foundation. The transmission electron microscopy was performed in the Microscopy Imaging Center (RRID SCR_018821) in the Larner College of Medicine at the University of Vermont. Excellent technical assistance at the Microscopy Imaging Center was provided by Michele von Turkovich and Tim Connolly. Ko-Wei Lui conducted the LIVE/DEAD BacLight Bacterial Viability Assay. The funders had no role in study design, data collection and analysis, decision to publish, or preparation of the manuscript.

## Author Contribution

Conceived and designed research: ZL, RB, AN, KK and BAS; performed experiments: ZL, RB, AN, CR, AA, SAG and DJT; ZL and TH analyzed data; all authors interpreted results of experiments; ZL and BAS drafted the manuscript: all authors edited and revised manuscript and approved the final version of the manuscript.

---

**Supplemental Figure 1.**
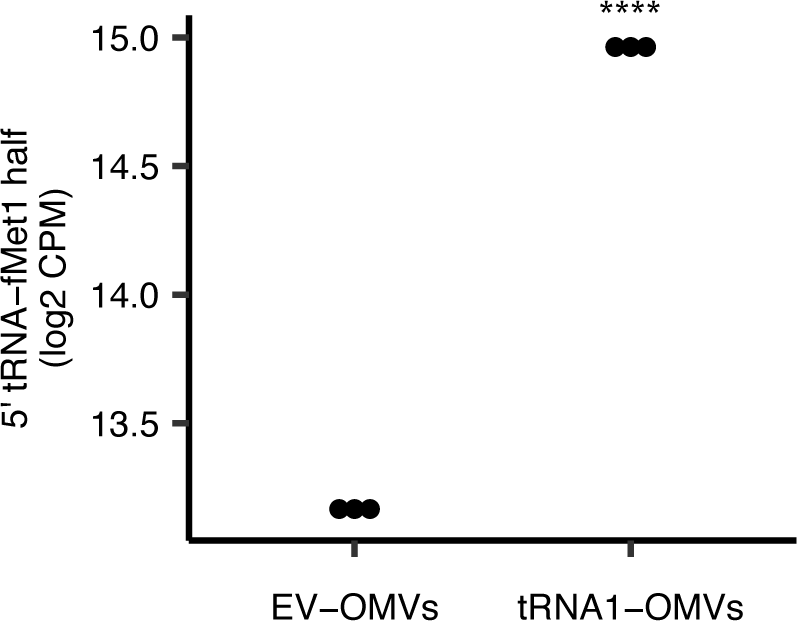
5′ tRNA-fMet1 half expression in tRNA1-OMVs compared to EV-OMVs. Small RNAs in EV-OMVs and tRNA1-OMVs harvested from LB culture with arabinose were subjected to small RNA sequencing to quantify 5′ tRNA-fMet1 half (n = 3). edgeR analysis (see methods) was used to determine significance. ****P<0.001

**Supplemental Table 1.**
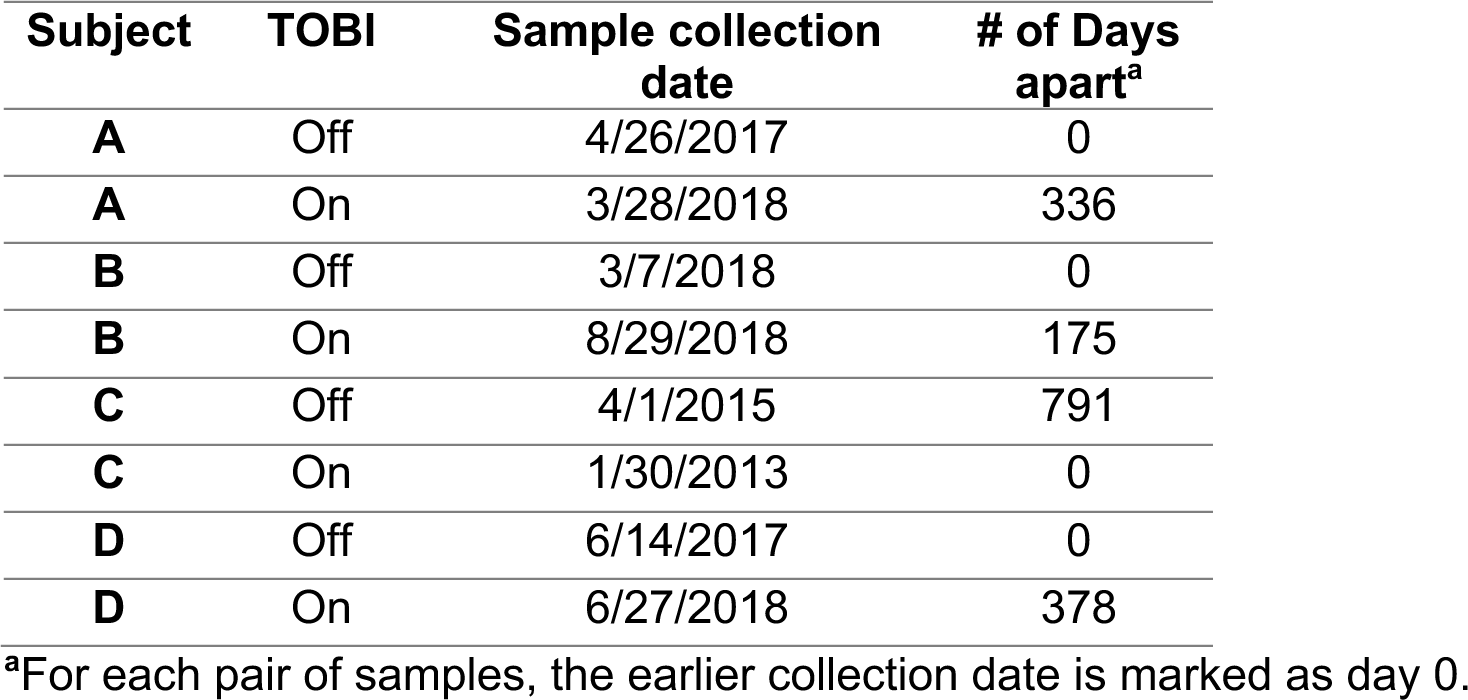
BALF sample collection dates.

| Subject | TOBI | Sample collection date | # of Days apart<sup>a</sup> |
| --- | --- | --- | --- |
| A | Off | 4/26/2017 | 0 |
| A | On | 3/28/2018 | 336 |
| B | Off | 3/7/2018 | 0 |
| B | On | 8/29/2018 | 175 |
| C | Off | 4/1/2015 | 791 |
| C | On | 1/30/2013 | 0 |
| D | Off | 6/14/2017 | 0 |
| D | On | 6/27/2018 | 378 |

<sup>a</sup>For each pair of samples, the earlier collection date is marked as day 0.

